# Genomic and evolutionary evidence for drought adaptation of grass allopolyploid Brachypodium hybridum

**DOI:** 10.1101/2024.10.06.616847

**Authors:** Yuanyuan Wang, Guang Chen, Fanrong Zeng, Fenglin Deng, Zujun Yang, Zhigang Han, Shengchun Xu, Eviatar Nevo, Pilar Catalán, Zhong-Hua Chen

## Abstract

Climate change increases the frequency and severity of drought worldwide, threatening the environmental resilience of cultivated grasses. However, the genetic diversity in many wild grasses could contribute to the development of climate-adapted varieties. Here, we elucidated the impact of polyploidy on drought response using allotetraploid *Brachypodium hybridum* (Bh) and its progenitor diploid species *B. stacei* (Bs). Our findings suggest that progenitor species’ genomic legacies resulting from hybridization and whole-genome duplications conferred greater ecological adaptive advantages to Bh over Bs. Genes related to stomatal regulation and immune response from S-subgenomes were under positive selection during speciation, underscoring their evolutionary importance in adapting to environmental stresses. Biased expression in polyploid subgenomes [*B. stacei*-type (Bhs) and *B. distachyon*-type (Bhd)] significantly influenced differential gene expression, with the dominant subgenome exhibiting more differential expression. *B. hybridum* adapted a drought escape strategy characterized by higher photosynthetic capacity and lower *WUEi* than Bs, driven by a highly correlated co-expression network involving genes in the circadian rhythm pathway. In summary, our study showed the influence of polyploidy on ecological and environmental adaptation and resilience in model *Brachypodium* grasses. These insights hold promise for informing the breeding of climate-resilient cereal crops and pasture grasses.

## Introduction

The global food production system needs to adapt to increasing events of abiotic stresses such as drought, heat, salinity and heavy metals (Adem *et al*., 2020; Bai *et al*., 2020; Lesk *et al*., 2016; Wang *et al*., 2023a). The majority of human calories are derived from grass species including staple cereal crops and animals fed with cereal grains and pasture grass (Godfray *et al*., 2010; Godfray *et al*., 2011). Therefore, global food security will benefit from a better understanding of abiotic stress resilience of grasses (Brkljacic *et al*., 2011). Since 1990s, *Brachypodium distachyon* has emerged as an ideal grass model (Brkljacic *et al*., 2011; Draper *et al*., 2001; Vogel *et al*., 2010; Decena et al. 2021; Hasterok et al. 2022; Minadakis et al. 2024) and its evolutionary relationship within the genus and with other grasses have been extensively studied (Catalán *et al*., 2016; Diaz-Perez et al. 2018; Gordon *et al*., 2020; Sancho et al. 2022). Recently, the polyploid *B. hybridum* and its two diploid progenitor species *B. distachyon* and *B. stacei* were selected as a model complex to investigate the multiple origins and the adaptive consequences of allopolyploidy (Gordon et al. 2020; Scarlett et al. 2022; Mu et al. 2023a, 2023b, Campos et al. 2024).

The annual *Brachypodium hybridum* is an allotetraploid (2n=4x=30; x=5+10) derived from the cross of *B. distachyon* (2n=2x=10; x=5) and *B. stacei* (2n=2x=20; x=10) diploid progenitor species followed by subsequent whole-genome duplications (WGD) (Catalán *et al*., 2012). Molecular evolutionary analysis indicated that *B. stacei* is the oldest diploid lineage within *Brachypodium*, splitting from the common ancestor ∼10 million years ago (Mya), followed by the divergence of the *B. distachyon* lineage at ∼7 Mya (Catalán *et al*., 2012; Sancho *et al*., 2018, 2022). In nature, plant diversity and speciation are often facilitated by recurrent interspecific hybridizations and allopolyploidizations (Ramsey, 1998) and *B. hybridum* formed at least three times from reciprocal crosses (Mu et al. 2023a). It was revealed that one D-plastotype (plastome derived from *B. distachyon*) and two S-plastotypes (plastome derived from *B. stacei*) lines were independently formed ∼1.4 Mya, and ∼0.14 Mya and ∼0.13 Mya, respectively (Gordon *et al*., 2020; Mu et al. 2023a). This evolutionary framework of ancestral and young allopolyploid plants provides unique opportunities for *Brachypodium* species to serve as bridges to accelerate the process of gene discovery for tolerance to abiotic and biotic stresses in cereal crops (Brutnell *et al*., 2015).

Ancient WGDs have occurred throughout the evolution of eukaryotes and a large proportion of major crops such as bread wheat, cotton, maize, soybean are polyploids (Masterson, 1994). The close relationship between stress and polyploidy has been manifested in the process of the establishment and development of polyploids. Polyploid plants usually have superior stress tolerance due to increased genetic variation and genomic complexity acquired through hybridization and polyploidy (Doyle and Coate, 2019; Eric Schranz *et al*., 2012; Soltis DE, 2014; Van de Peer *et al*., 2017). WGD may have provide better chances for terrestrial adaptation of plants in the stressful transition from water to land (Liu *et al*., 2020). It was reported that ancient WGDs overlap with periods of dramatic global changes (Fawcett *et al*., 2009; Landis *et al*., 2018; Van de Peer *et al*., 2020; Vanneste *et al*., 2014), particularly in the adaptation to extreme environments (Bromham *et al*., 2020; Cai *et al*., 2020; Caperta *et al*., 2020; Deng *et al*., 2012; Manzaneda *et al*., 2012; van Laere *et al*., 2011; Zhang *et al*., 2020). The narrow genetic variation in many cultivated crops has restrained the development of climate-adapted varieties. Fortunately, their wild relatives provide a vast and largely untapped reservoir of genetic variation for crop improvement (King *et al*., 2013) because some stress tolerance traits of wild species have not been altered by human selection during crop domestication in the last 10,000 years. Polyploidy-triggered gene duplication is vital for crop domestication and breeding for stress resistance (Renny-Byfield and Wendel, 2014). For example, in some angiosperm groups polyploids occupy drier habitats than their closely related diploids (Gunn *et al*., 2020; Te Beest *et al*., 2012), like autotetraploid *Arabidopsis* (Chao *et al*., 2013), rice (*Oryza sativa*) (Yang *et al*., 2014b), watermelon (*Citrullus lanatus*) (Zhu *et al*., 2018), and allotetraploid citrange (*Citrus sinensis* × *Poncirus trifoliata*) (Ruiz *et al*., 2016) and allohexaploid wheat (*Triticum. aestivum* L., AABBDD) (Yang *et al*., 2014a). Understanding the role of polyploidy in *Brachypodium* to adapt to environmental challenges can provide important insights into the breeding for climate-adapted cereal crops. However, the mechanistic basis of intraspecific and intra(sub)genomic variation of *Brachypodium*, especially the relation between drought tolerance and ploidy segregation is still elusive.

Opposite slopes at Mount Carmel of Israel designated as ’Evolution Canyon’ (EC) display large microclimatic contrasts between the hotter and drier tropical African South-facing Slope (AS) and the temperate European North-facing Slope (ES) (Kang *et al*., 2019; Li *et al*., 2016; Nevo, 1995; Wang *et al*., 2020a; Yablonovitch *et al*., 2017). The EC model reveals evolution at a microscale involving biodiversity divergence, adaptation, and incipient sympatric speciation across life from viruses and bacteria through fungi, plants, and animals (Nevo, 2012). In our previous study, we employed wild barley, wild emmer wheat, *B. stacei*, and *B. hybridum*, from EC to investigate the role of diploid and polyploid differentiation in the adaptation of grasses to drought (Wang *et al*., 2023b). The current study aims to elucidate the role of polyploidy in drought response strategies by comparing the physiological traits, genomic variation, and biased homeologs’ expression between the *B. stacei* genome (Bs) and the two sub-genomes of *B. hybridum* (Bhs, Bhd).

## Results

### Polyploidy and Genomic adaptation of B. hybridum and B. stacei

Previously, we identified 19 *Brachypodium* accessions from AS and ES using centromeric and telomeric-specific probes in fluorescence *in situ* hybridization (FISH) analysis, of which *B. hybridum* (Bh) individuals were mainly from the dry and high light AS and *B. stacei* (Bs) individuals from the wet and shady ES (Wang et al., 2023b). The more mesic progenitor species *B. distachyon* is not present today in EC (Mu et al. 2023a). The time of divergence of genomes and/or subgenomes pairs can be estimated from the synonymous substitution rates (Ks) using the equation T (the divergence time) = Ks/2λ, where λ = 6.5 × 10^−9^ (He *et al*., 2013; Lynch and Conery, 2000; Takahagi *et al*., 2018) is the absolute mean rate of substitutions per site per year inferred for the grass genes (Fig. 1A). The Circos plot of intergenomic collinearity showed highly conserved blocks of genomic synteny of *B. stacei* and BhS (Fig. 1B, Table S1). The *B. hybridum* genome consists of a substantial amount of long terminal repeats (LTRs) such as Copia and Gypsy in the centromeric regions of the BhS and BhD chromosomes (Fig. 1B, Table S2). The BhD subgenome has 31% more Gypsy than the BhS subgenome per kb, while the BhS subgenome has 25% more Gypsy and 32% more Copia than the Bs diploid genome.

**Figure 1.**
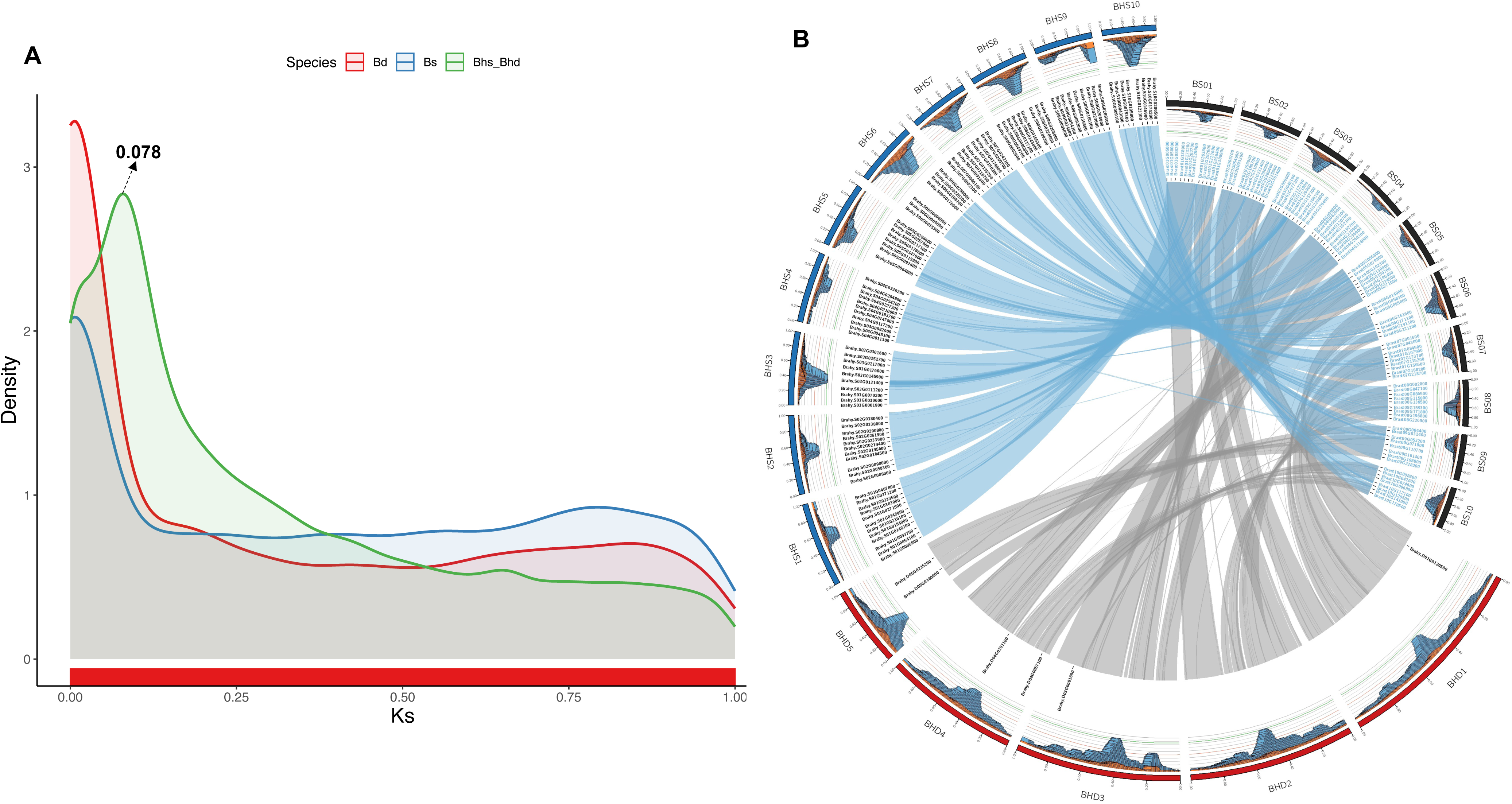
Genomic evolution analysis of *Brachypodium* species. (A) Ks density of *Brachypodium hybridum* (Bh), *Brachypodium stacei* (Bs), *Brachypodium distachyon* (Bd), BhD versus Bs (Bhd_Bs), and BhS versus Bd (Bhs_Bd). Ks values >1 were removed to eliminate saturated synonymous sites. The three peaks represent the divergence time between the sub-genomes of the allotetraploid and each diploid genome and WGD events of the allotetraploid. (B) Circos diagram depicting genomic relationships of Bh and Bs. The chromosomes are shown along with their relative position. The outer track shows the distribution of LTR retrotransposons over each chromosome as stacked histograms, blue bars and orange bars represent Copia-domain distribution and Gypsy-domain distribution, respectively. Text of inner ring demonstrates homeolog genes with highest Ka/Ks values between the two species. The inner arcs designate inter-genomic rearrangements.

To investigate the genes under positive selection during the evolution of *B. hybridum*, we calculated the Ka/Ks ratio for ortho-homeologs between *B. hybridum* and *B. stacei*. Functional enrichment analysis using Gene Ontology (GO) revealed that genes with a Ka/Ks ratio greater than 1 are significantly involved in processes such as stomatal movement, stomatal complex development, immune response, and transmembrane transport (Fig. S1). These functions demonstrate the evolutionary importance of these genes in adapting to environmental stresses. Therefore, we compared the stomatal traits between Bh and Bs (Fig. 2A-E). Under well-watered conditions, Bh exhibited significantly greater aperture length, guard cell length and width, and subsidiary cell length compared to Bs, which could be due to the polyploidy (higher number of gene copies) of Bh resulting in bigger cell size. Bs displayed narrower aperture width and subsidiary cell width. Drought stress led to a reduction in stomatal length and width in both *Brachypodium* species, with a more pronounced reduction in stomatal width in Bs (Fig. 2E).

**Figure 2.**
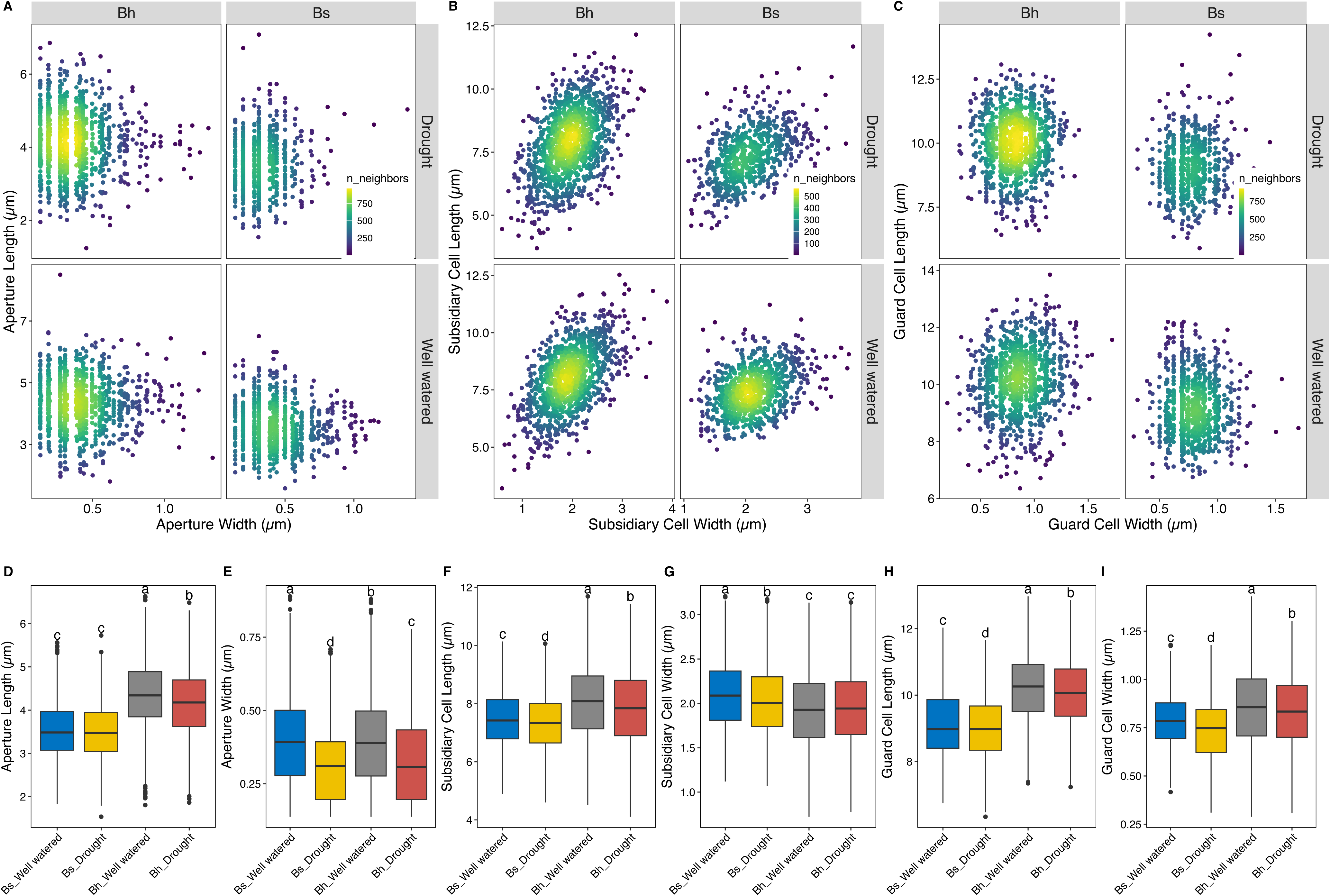
Stomatal structure of *B. stacei* and *B. hybridum* in response to drought. Stomatal aperture (A), subsidiary cell (B), guard cell (C), comparison of stomatal parameters between *Brachypodium stacei* and *Brachypodium hybridum.* The data are visualized using scatter density plots, where the color gradient represents the density of neighboring data points (n.neighbors), with yellow indicating higher density and purple indicating lower density. Top panels show measurements under drought conditions, while bottom panels show measurements under well-watered conditions. The results highlight differences in stomatal and subsidiary cell structure between the two species and the impact of drought stress on these parameters. Differences between groups were determined using one-way ANOVA with a post-hoc Tukey HSD test. Differences were considered significant when *p* < 0.05. (D-I).

### The transcriptional landscapes of drought response in B. hybridum and B. stacei

Bh and Bs accessions showed distinct stomatal and photosynthetic responses to drought. Under drought treatment, Bh accessions maintained significantly higher photosynthesis rate, stomatal conductance, stomatal aperture area, lower water use efficiency (*WUEi*) (Figs. S2) and had earlier flowering time than *B. stacei* (Wang *et al*., 2023b). To reveal the molecular mechanisms underlying the physiological difference between Bh and Bs, we conducted transcriptome experiments of these *Brachypodium* accessions under drought treatment (Table S3). The transcriptomic profiles between Bs and Bh, Bs and Bh’s sub-genomes can be clearly separated by PCA (Fig. S3). We identified ortho-homeolog triads using the collinear results for a global comparison between the Bs genome and the BhS and BhD subgenomes of *B. hybridum*. Genes differentially expressed in more than 30% (i.e. >3) accessions were identified as differentially expressed genes (DEGs) in this species, Venn plot showed Bs share more DEGs with BhS than with BhD (Fig. 3A). Shared DEGs in BhS, BhD and Bs were primarily involved in responses to environmental stress, such as light intensity, starvation, nutrient level and hypoxia, while unique DEGs in BhS showed enrichment in circadian rhythm and rhythmic process, and in Bs were associated with pollen recognition and defense responses (Fig 3B).

**Figure 3.**
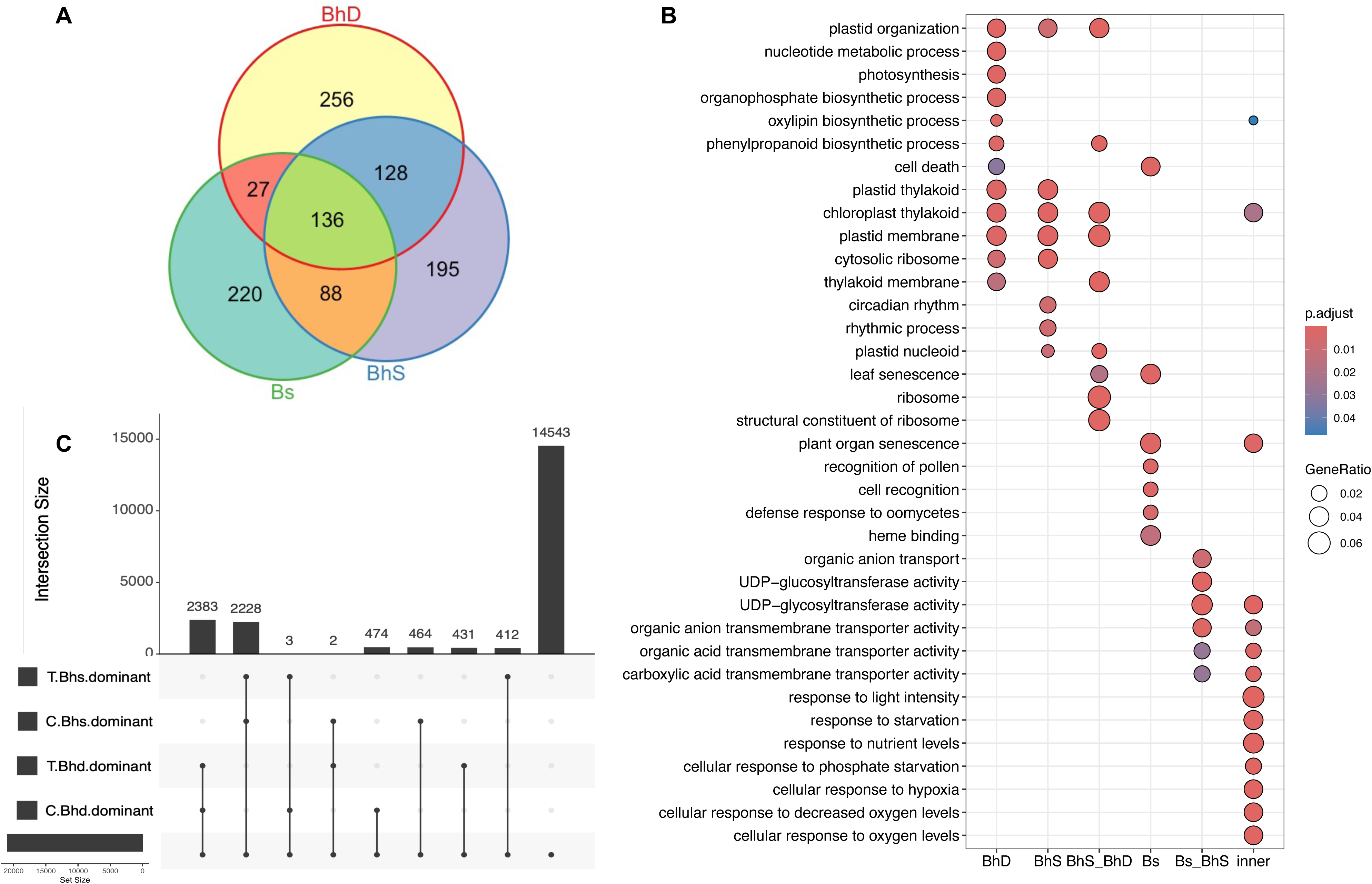
Global transcriptome analyses of drought response in *B. hybridum* and *B. stacei*. (A) Venn plot for DEGs in sub-genomes of *B. hybridum* and in *B. stacei,* genes that are differentially expressed in more than 30% of the accessions within each species were considered DEGs. (B) Gene Ontology (GO) enrichment analysis for DEGs intersections in Venn plot, ‘inner’ represent the 136 genes differentially expressed in Bhd, Bhs and Bs. (C) Expression differentiation of paired homeolog genes between *B. hybridum* subgenomes. Homeolog gene pairs were identified by collinear blocks using the Bhs subgenome as reference for each pair. The black horizontal bars on the left of the figure show the number of paired homeolog genes dominated by Bhs/Bhd subgenome under well water (Control, C) or drought (Treatment, T). The lined dots indicate genes share the same biased homeolog expression pattern. The gene numbers of intersections are represented by vertical bars.

We further assessed biased expression pattern under well-watered condition and drought to validate whether drought governs shifts in gene expression between BhS and BhD. Although most genes are balanced expressed, there were 2,383 and 2,228 pairs of ortho-homeologs dominated by BhS or BhD under both the control and drought condition, (Fig. 3C), these two sets of genes are enriched in different biological processes. BhS genes are primarily enriched in functions related to environmental stimuli, such as defense response to bacterium and response to heat (Fig. S4). In contrast, BhD genes have more DEGs related to DNA damage stimulus, DNA repair, and regulation of the cell cycle (Fig. S4). Moreover, 5 ortho-homeologs changed their dominant sub-genome after drought treatment (Table S4). We compared the proportion of balanced versus unbalanced expression patterns in non-DEGs (Fig. 4A) and DEGs (Fig. 4B) across species/subgenomes. The results showed that 70% of non-DEGs exhibited balanced expression, while this proportion dropped to 42%-49% among DEGs. Notably, among DEGs in the BhD subgenome and/or Bs, BhD-dominant genes constitute the largest proportion. Conversely, among DEGs in the BhS subgenome and/or Bs, BhS-dominant genes make up the greater proportion but not as high as in the previous case? (Fig. 4B). This indicates that biased expression in the subgenomes of polyploid Bh is significantly influenced by parental legacy, with the dominant subgenome (BhD) being more prone to differential expression among DEGs induced by drought.

**Figure 4.**
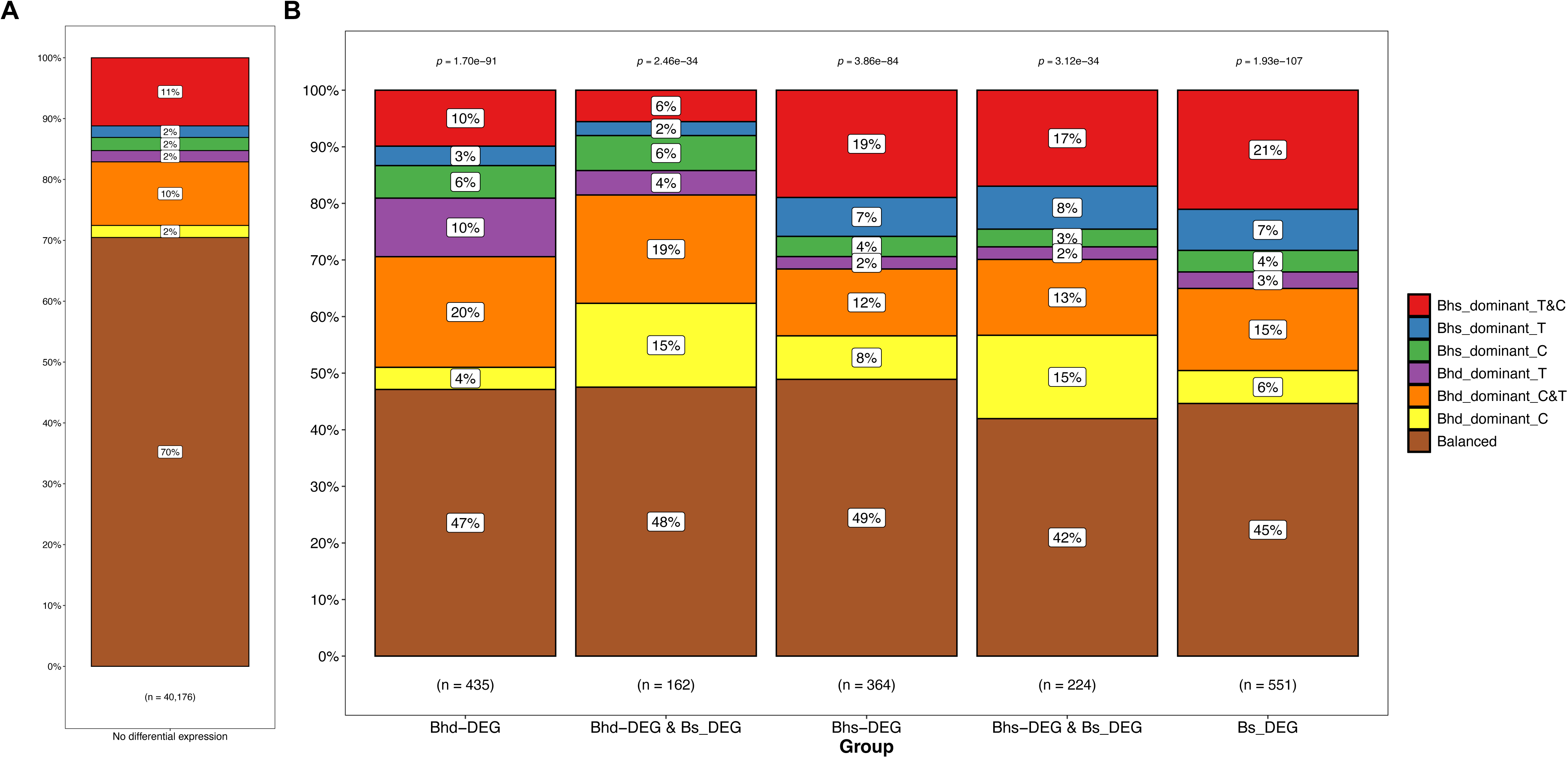
The proportion of non-differentially expressed genes (A) and DEGs (B) balanced or predominantly expressed in each sub-genome of *B. hybridum*. The different colors distinguish the sub-genomes that dominate gene expression under control (well-watered) or treatment (drought) conditions. Each bar represents a specific category of DEGs, with “n” shows the number of genes in each category. The p-values displayed above each bar denote the statistical relevance among different expression patterns in each category.

We employed RNAseq and physiological traits of each accession to conduct weighted correlation network analysis (WGCNA). Overall, 24 and 20 co-expression gene modules were clustered in Bh and Bs, respectively. The correlations among physiological traits were similar in Bh and Bs (Fig. 5). Moreover, Bh showed more co-expression modules significantly correlated with the physiological traits. Notably, ortho-homeologs in Bh that are enriched in ‘circadian rhythm’, ‘rhythmic process’, and ‘response to blue light’ showed a positive correlation with net photosynthetic rate (*A*) and stomatal conductance (*gs*), but a negative correlation with *WUEi* (Fig. 6). In the co-expression network, several highly connected hub genes were identified, many of which are enriched in Mitogen-Activated Protein Kinase (MAPK) Kyoto Encyclopedia of Genes and Genomes (KEGG) pathway (Figure S5). An important difference between Bh and Bs in MAPK signaling was the regulation of genes encoding the plasma membrane receptor kinase FLAGELLIN SENSING 2 (FLS2), which participates in pathogen attack triggered stomatal closure (Spallek *et al*., 2013). The expression of genes involved in the ABA signaling pathway (*OST1*, *open stomata 1*; *HAB*, *Hypersensitive to ABA; PYL5*, *pyrabactin resistance 1-like 5*; *HAI, highly ABA-induced PP2C gene 1; CBL-interacting protein kinase)* are very similar between Bh and Bs. It thus indicates that although drought primarily affected ABA signalling pathway, it also regulated other networks particularly involved in the key hubs like MAPK (Figures S6, S7).

**Figure 5.**
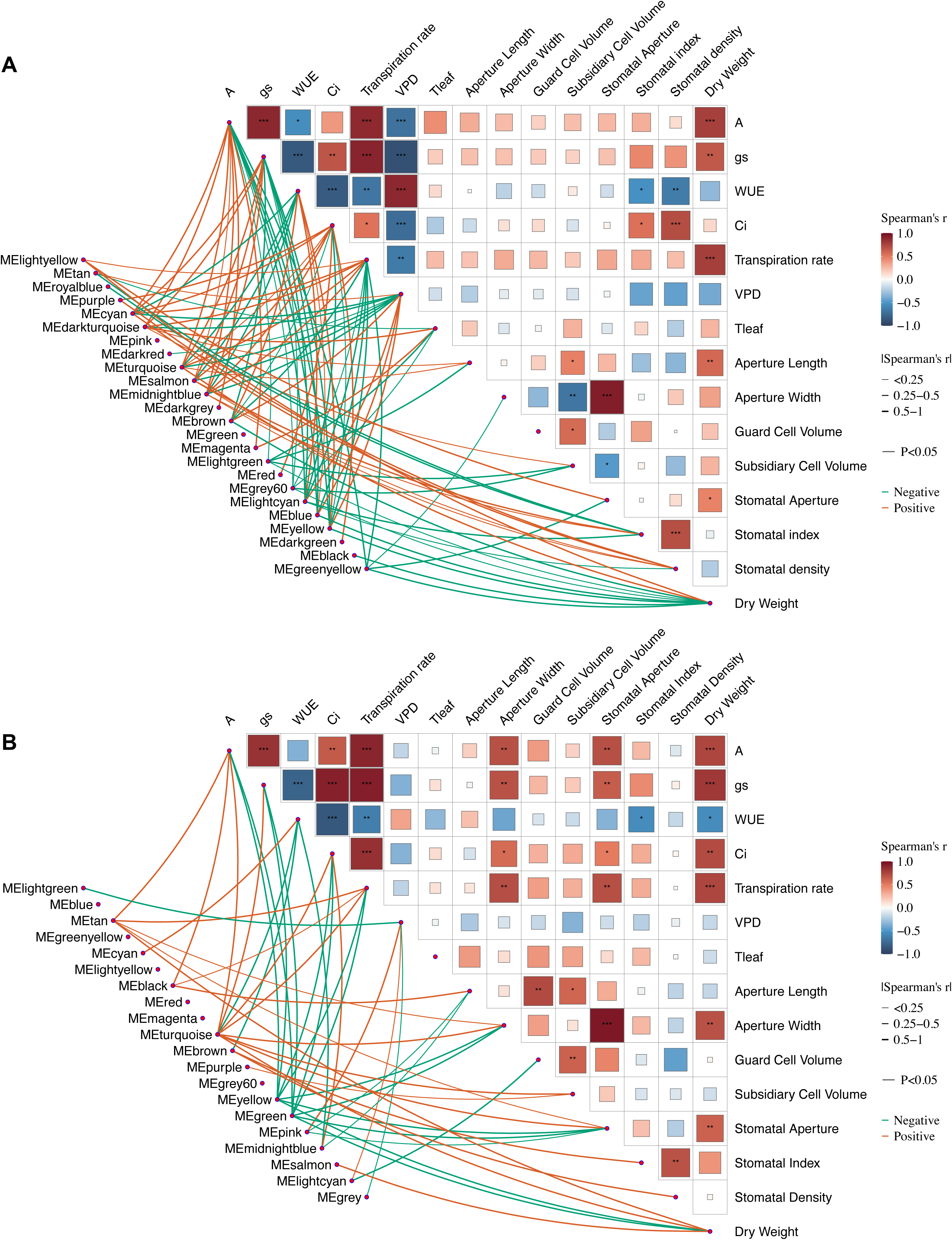
Significant associations between gene clusters and physiological traits in (A) *B. hybridum* and (B) *B. stacei.* A triangular correlation plot was conducted to display the relationships between physiological or stomatal parameters. Red indicates positive correlations, while blue represents negative correlations, asterisks denote the significance of the correlations. The lines connected to each parameter represent co-expressed gene modules, with only those showing significant correlations retained. Red lines indicate positive correlations, green lines indicate negative correlations, and the thickness of the lines represents the Spearman’s r value.

**Figure 6.**
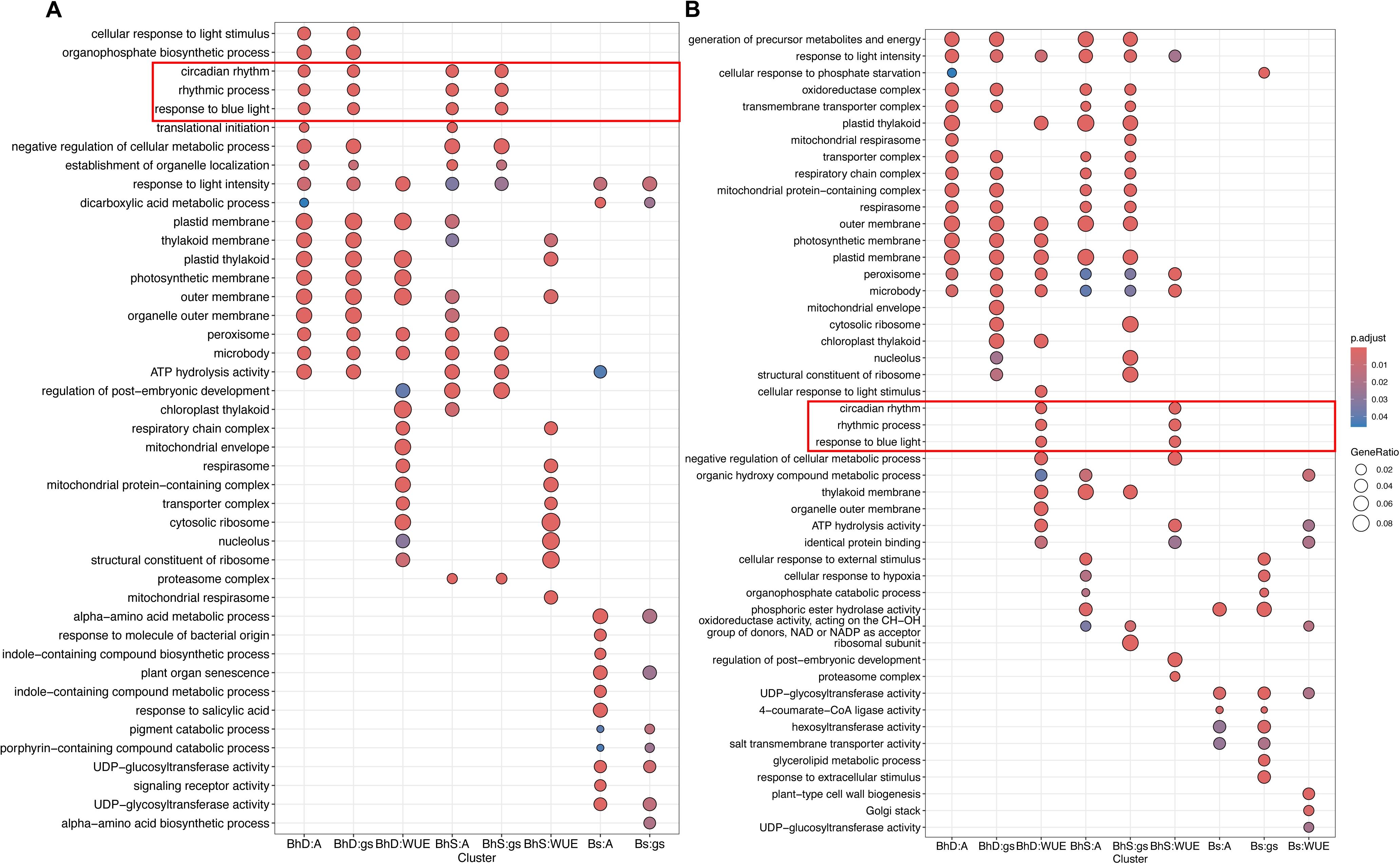
GO terms of gene modules significantly positively (A) and negatively (B) correlated with physiological traits across different subgenomes.

### Genomic variants and positive selections in B. hybridum and B. stacei

To identify potential adaptive alleles in response to drought environment in the AS, genomes of all 19 accessions of Bh and Bs were re-sequenced, yielding 794 Gb of clean data (Table S3). After filtering, 1,142,287 and 614,074 high-confidence SNPs, along with 165,880 and 99,276 indels, were identified in Bh and Bs, respectively. These variants were distributed across 2,090 genes in Bh and 669 genes in Bs (Fig. 7A). Among these, 363 and 140 genes in Bh and Bs, respectively, contained high-impact variants, with annotation for their Arabidopsis homologs detailed in supplementary Table S5. Notably, some genes overlapped between Bh and Bs, particularly those with NB-ARC and P450 domains or those involved in plant resistance and plant-pathogen or herbivore interactions. Based on piN and piS calculated from SNPs, genes under positive selection (piN/piS > 1) in the two subgenomes of Bh (Table S6) were primarily associated with stomatal movement regulation and immune response. However, they were linked to nitrate response, immune response, biosynthetic processes, and calcium binding in Bs (Fig. 7B). DEGs of both subgenomes Bs and Bd under similar positive selection constrains werealso similarly expressed (Table S6; Figs S9-S10). Interestingly, these positively selected genes were also influenced by biased ortho-homeolog expression. For example, in BhS, genes under positive selection showed a reduction in the proportion of BhD-dominant homeologs, while in BhD genes under positive selection exhibited a decreased proportion of BhS-dominant homeologs (Fig. 7C).

**Figure 7.**
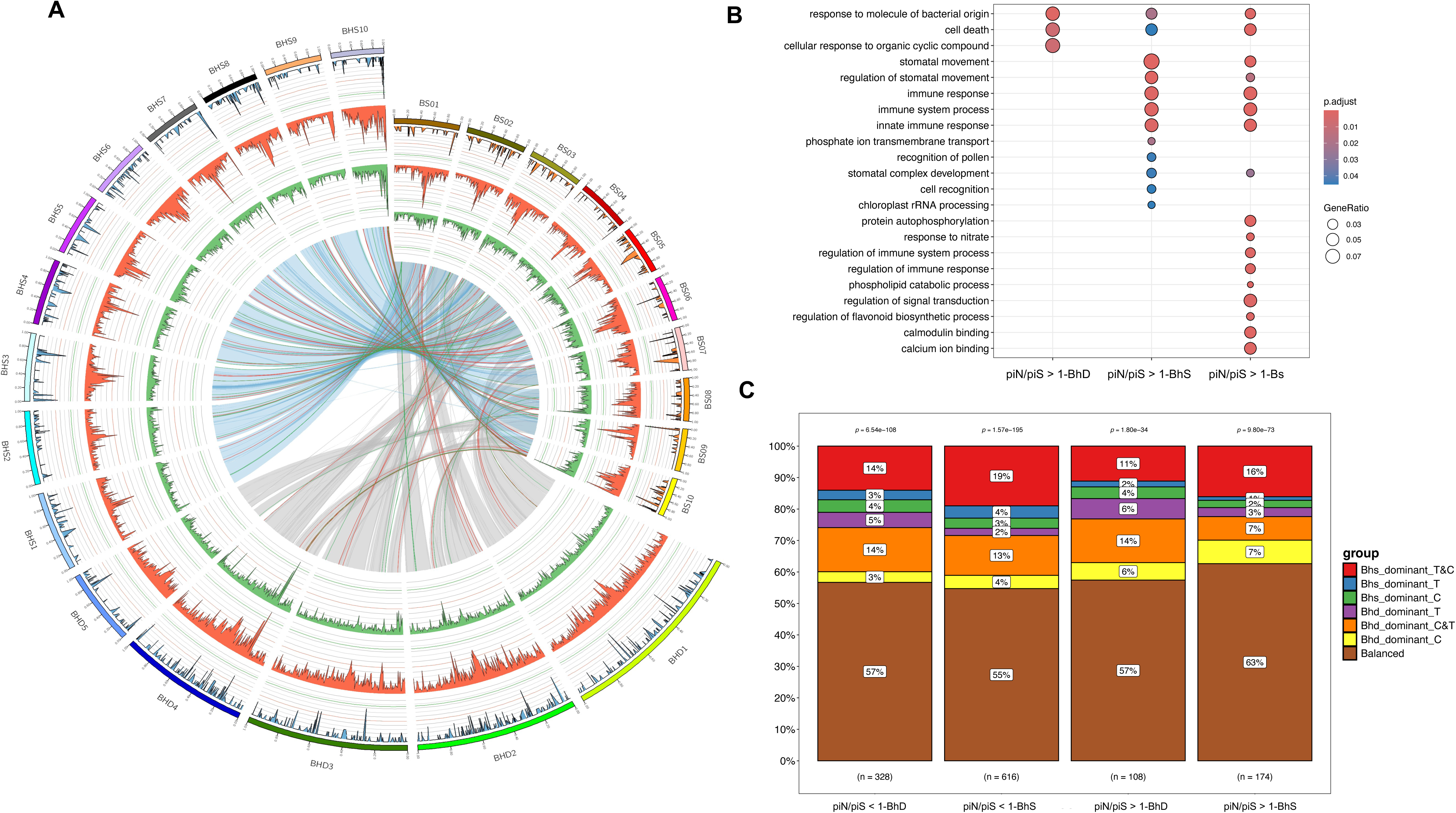
Genome resequencing and the relationship between genes under selective pressure and expression patterns in *B. hybridum and B. stacei*. (A) Circos plot representing from outer to inner circles the number of variants with high impact (gain/lost stop_codon) of each gene of Bh (blue) and Bs (orange), single nucleotide polymorphism (SNP) density (red) and Indel density (green). The inner arcs designate inter-chromosomal rearrangements, red and green lines represent enriched high impact variants of Bh or Bs, respectively. (B) GO terms for genes with high impact SNP. (B) piN/piS indicates ratio of nonsynonymous to synonymous SNP. (C) The proportion of genes under selective pressures with balanced or predominantly expressed in each sub-genome of *B. hybridum*. The different colors distinguish the sub-genomes that show dominant gene expression under control (well-watered) or treatment (drought) conditions. Each bar represents a specific category of genes, with “n” showing the number of genes in each category. The p-values displayed above each bar denote the statistical relevance among different expression patterns in each category.

## Discussion

### Recent hybridization and WGDs provide drought adaptive potential for *B. hybridum*

Numerous studies emphasized the impact of hybridization and WGD on gene expression pattern (Feldman *et al*., 2012; Ramírez-González *et al*., 2018; Renny-Byfield and Wendel, 2014) through neofunctionalization of duplicated genes (Kondrashov, 2012) or tissue-specific expression of genes (Makova and Li, 2003). In general, polyploids can display rapid responses under stress. Previous studies in cotton (Jiang *et al*., 1998), wheat (Krasileva *et al*., 2017; Ramírez-González *et al*., 2018), maize (Goff, 2011), and *Cucurbita* (Sun *et al*., 2017) revealed that increased genetic diversity triggers superiority of polyploids under abiotic stresses. Here, we found that a subset of genes of Bh was dominated by a single subgenome in both well-water and drought conditions, thus, the transcriptional regulation of allotetraploid *B. hybridum* is shaped by subgenome-dominant expression patterns. Furthermore, we also validated the hypothesis that allopolyploid species utilize the expression patterns of both progenitor subgenomes depending on environments (Table S4) (Shimizu-Inatsugi *et al*., 2017).

Collinearity of genes in plant genomes generally decreases with increasing evolutionary distance (Wicker *et al*., 2010). Here we showed that the two subgenomes of *B. hybridum* share strongly conserved syntenic blocks with their corresponding progenitor diploids (*B. stacei*, Figs. 1 and 7; *B. distachyon*, Gordon et al. 2020). It was reported that syntenic break points (SBPs) are hotspots of evolutionary novelty that could act as a reservoir for producing adaptive phenotypes (Murat *et al*., 2010). The deranged blocks may be disturbed by chromosomal recombination during interspecific hybridization. However, strong karyotypic differences between the Bs and Bd chromosomes probably prevented intersubgenomic chromosome rearrangements in *B. hybridum* (Mu et al. 2023a). The evolutionary potential and high frequency of polyploidy or hybridization in flowering plants are also due to the activity of transposable elements (TEs) among which LTRs are abundant (Oliver *et al*., 2013). They have strong impact on the structure, function, evolution, and especially genome size of their host genome (Baidouri and Panaud, 2013). Here, the density distribution of Gypsy-type LTRs in Bh is obliviously higher than in Bs, it is because *B. hybridum* shows an additive pattern in LTRs; its LTR content is the results of the sum of LTRs from both progenitor species (BhS and BhD subgenomes). However, the proportion with respect to the respective genome size is quite similar (Mu et al., 2023a).

### The crucial role of circadian rhythm regulation in B. hybridum sub-genome for its escape from drought stress

Exploration of divergent drought tolerance strategies between tetraploids and their diploid progenitor species will provide new insight on crop breeding with improved adaptation to unpredictable environments. Under drought condition, plants inevitably have a trade-off between carbon accumulation and the risk of deleterious soil water depletion, hence plants have developed several strategies to cope with drought stress, including dehydration avoidance, drought tolerance, and drought escape (McKay *et al*., 2003; Sherrard and Maherali, 2006; Tuberosa, 2012). The underlying mechanism of the drought escape strategy employed by *B. hybridum* is likely to be regulated by light perception and circadian clock to promote early flowering (Du *et al*., 2018). Also, early flowering can be achieved by maintaining rapid growth and high stomatal opening and photosynthesis, allowing the plant to complete its life cycle before the onset of severe water stress (Donovan *et al*., 2007; Heschel and Riginos, 2005; Ivey and Carr, 2012; Manzaneda *et al*., 2015; Sherrard and Maherali, 2006; Wu *et al*., 2010).

Stomata exert extensive influences on plant growth. Stomatal function is constrained by a safety-efficiency trade-off because the survival rate of plants depends on stomatal closure under drought (Henry *et al*., 2019). It has been reported that stomatal size varies with WUE, facilitating plant adaptive plasticity under drought (Dittberner *et al*., 2018). Stomatal guard cell length was reported to be a useful trait to separate *B. distachyon* complex species and other related diploid and polyploid grasses (Borrino and Powell, 1988; Catalán *et al*., 2012; Katsiotis and Forsberg, 1995). We found that *B. hybridum* accessions has larger stomata size than *B. stacei* (Fig. 2), concurring with similar results when the allotetraploid is compared with its alternative progenitor species *B. distachyon* (Catalán et al. 2012; Manzaneda et al. 2015). The suite of specialized stomatal structures altered by polyploidy of the annual *Brachypodium* species may result in greater capacity of *B. hybridum* in drought tolerance.

The internal timekeeper circadian clock of plants can anticipate environment stimuli such as light and temperature to prepare for upcoming challenges. The plant circadian clock also participates in multiple regulatory pathways such as photosynthesis, seed germination, stomatal movement, pollination, flowering and senescence (Srivastava *et al*., 2019). Therefore, we propose that the interconnection between ABA signaling, and circadian rhythmic regulation may explain why drought-escapist *B. hybridum* accessions maintain higher stomatal conductance and stomatal aperture area (Figures 5-6), implying that circadian clock provides optimal utilization of machinery to obtain improved adaptation during abiotic stresses (Grundy *et al*., 2015).

### Linking drought stress signalling with biotic resistance in Brachypodium stacei

Water deficit affects a large spectrum of plant functions such as transpiration, photosynthesis, tissue growth, and reproductive development (Chaves *et al*., 2003; Khalid *et al*., 2023). Components of MAPK cascades are regarded as converging points of abiotic as well as biotic stress signaling (Chinnusamy *et al*., 2004; Huang *et al*., 2012; Xiong and Yang, 2003), which are activated by the PYR/PYL/RCAR-SnRK2-PP2C ABA signaling modules (Danquah *et al*., 2015). In our study, MAPK pathway connects the drought-induced ABA signaling to plant immune receptor FLS2 (Fig. S6), which recognizes bacterial flagellin and its immunogenic epitope flg22 to inhibit infection by bacterial pathogens (Wang *et al*., 2020b). Interestingly, it was reported that MAPK activation links the late endocytic trafficking of FLS2 specifically with defense-associated stomatal closure (Spallek *et al*., 2013). We found that the FLS2 and its recognition receptors (Spallek *et al*., 2013) were up-regulated in *B.stacei* but down-regulated in *B.hybridum* (Fig. S7?), unveiling that drought-induced stomatal closure not only affects the expression of genes involved in drought regulation, but also influences other pathways. Our results highlight the significance of the immune related response in the evolution of *Brachypodium* genotypes (Figs 7B; S1; S8), implying that the diploid progenitor *B. stacei* genome may have dominated the response to biotic stresses.

### Implications of B. hybridum and B. stacei in stress tolerance of grasses

Adaptation is an evolutionary process where traits in a population are naturally selected to better meet the environmental challenges (Singaravelan *et al*., 2008). It was found that the natural habitats for *B. hybridum* overlap with but are even broader than those of its diploid progenitor species (Catalán *et al*., 2016; López-Alvarez *et al*., 2015). In our study, we selected *B. hybridum* individuals from the dry and high light AS, the only slope in Evolution Canyon where it grows (Mu et al. 2023a), and *B. stacei* individuals from the forested and shady ES. However, it is of importance to note that *B.stacei* also grows in AS, typically sheltered under shrubs (Mu et al. 2023b), while *B.hybridum* is a heliophyte. The AS receives 200-800% higher solar radiation than the ES, where high light, heat, and drought are the major environmental constraints on the AS, while shade and pathogen are the major constraints on the ES (Nevo, 2012). The inter-slope differentiation pattern is shaped by ecological factors, for instance, the abundance of ‘water-dependent’ organismal groups is significantly lower on AS than in ES (Melamud *et al*., 2007), while the diversity in soil microfungal community structure was related to variations in soil moisture and solar radiation (Grishkan and Nevo, 2008). *B. stacei* usually lives in shady habitats, like the understories of pine forests, juniper woods, and inside spiny shrubs, which protect the plant from direct insolation (Catalán et al. 2016b). By contrast, the ecologically ample *B. hybridum* successfully diversifies in both mesic and arid ecosystems (Lopez-Alvarez et al. 2015), mostly grows in sunny open habitats (Catalán et al. 2016b). Thus, the two *Brachypodium* species exhibit unique advantages towards ecological and environmental adaptation to guide breeding efforts on climate resilient cereal crops and pasture grass in the future.

## Materials and methods

### Plant materials and growth condition

Accessions were collected from Lower Nahal Oren, Mount Carmel, Israel, at the ‘Evolution Canyon’ African Slope (AS) and European Slope (ES). Plant growth condition followed those described in (Chen *et al*., 2019). Plants were cultivated with the same soil and fertilizer condition at 22 ± 2 °C (day) and 20 ± 2 °C (night) and 60% relative humidity (RH) under a 14h/10h light/dark photoperiod in a growth chamber. The drought treatments were induced three weeks after germination and monitored by PR2 multi-depth soil moisture probes (Dynamax, USA). Gas exchange measurement and stomatal assay were taken and processed when the water holding capacity was gradually maintained at 10% (v/v) in drought-treated plants. Leaf samples of well-watered and drought-treated plants were collected for RNA sequencing at the same time. At least three biological replicates were used for each sample and treatment.

### Gas exchange measurement and stomatal assay

Net CO_2_ assimilation (*A*), stomatal conductance (*g_s_*), intercellular CO_2_ concentration (*Ci*), transpiration rate (*T_r_*), leaf vapor pressure deficit (VPD) and leaf temperature (T_leaf_) were measured with LI-6400 infrared gas analyzer (Li-Cor Inc., Lincoln, NE, USA) (Chen *et al*., 2019; Liu *et al*., 2017). The measured parameters were set at flow rate: 500 mol s^−1^, saturating PAR: 1500 mol m^−2^ s^−1^, 400 mmol mol^−1^ CO_2_, relative humidity: 60%.

The stomatal assay was performed on abaxial epidermises and measurements were carried out as described previously (Chen *et al*., 2019; Liu *et al*., 2017). The stomatal assay was conducted from 10 am to 4 pm in parallel with the gas exchange measurement. Abaxial epidermises were peeled and immersed in the measuring solution (10 mM KCl, 5 mM Ca^2+^-MES, pH 6.1) and mounted on slides. For each sample, at least 30 images were taken from at least five epidermis strips using a microscope with a CCD camera (Nikon, Tokyo, Japan). Stomatal parameters including aperture length and width, aperture width/ length, stomatal pore area, guard cell length, width and volume, subsidiary cell length, width and volume, stomatal density and index were measured by Image J software (NIH, USA). The ratio between net CO_2_ assimilation rate and stomatal conductance was calculated as intrinsic water-use efficiency (*WUE_i_*).

### Genome Evolution Analysis

The command line program wgd and related tools including BLASTP, MCL, PAML4, MAFFT, FastTree, I-ADHoRe were used for WGD analysis (Zwaenepoel and Van De Peer, 2019). CDS files used in WGD analysis from three species of the complex were downloaded from Phytozome (https://phytozome-next.jgi.doe.gov/info/Bstacei_v1_1; https://phytozome-next.jgi.doe.gov/info/Bhybridum_v1_1; https://phytozome-next.jgi.doe.gov/info/Bdistachyon_v3_1), codeml of PAML was used for calculating Ka/Ks. Syntenic homeologs were identified for each pairwise species comparison with the MCScanX algorithm (Wang *et al*., 2012) using default parameters. To obtain the collinearity both between and within species, an all-to-all BLASTP (version 2.9.0). The top 5 hits of each query were outputted and the final collinearity result was accessed by MCScanX. LTR-RTs (LTR retrotransposons) were identified using the LTR retriever (version 2.8) (Ou and Jiang, 2018). LTR_FINDER was used to detect full-length LTR-RTs in CDS sequences in *B. hybridum* and *B. stacei* genomes.

### RNA-sequencing and data analysis

The RNA library preparation and the alignment for sequencing reads were conducted in previous study (Wang *et al*., 2023b). the dominant sub-genome identification was based on *B. hybridum* homeolog gene expression characterized through the syntenic blocks. DESeq2 (Love *et al*., 2014) was used to assess biased Bhs *vs* Bhd homeologs expression patterns. For well-watered control (C) and drought treatment (T) conditions, each pair of homeologs was split into two groups; homeologs in Bhs were set as the reference group. For a given gene, if the expression of the Bhd homeolog was significantly higher than that of Bhs homeolog, it was designated ‘Bhd-dominant’, and the reverse was designated ‘Bhs-dominant’, while homeologs showing equal expression were categorized as ‘not biased’. The comparison of gene numbers showing biased and unbiased homeolog expression within or among the genotypes were visualized using UpSetR (R package).

WGCNA (Langfelder, 2008) was used to find clusters and modules of highly correlated co-expressed genes of *B. hybridum* and *B. stacei.* GO and KEGG enrichment were created using clusterProfiler (R package). The genic compositions of *B. hybridum* and *B.stacei* were determined with BLASTP using as reference the gene annotation database of *Arabidopsis thaliana* (org.At.tair.db). Initial components of protein-to-protein interaction networks were retrieved from the identified hub genes of the MAPK/circadian rhythm KEGG pathway. Public annotation data from Cytoscape (Su *et al*., 2014) was used for the interaction network construction. In order to use the familiar gene name instead of the name automatically generated by the software, network node and edge files were exported, then transferred (the ENTREZID) into SYMBOL of *Arabidopsis thaliana* (clusterProfiler, R package). The new node and edge files were imported into the software again, and the DEGs column was used as the group information to label color, whereas mapping type was done by continuous mapping.

### Genome resequencing and data analysis

The DNA isolation and library construction were conducted according to published work (Wang et al., 2023b). Sequence variants were discovered and base quality recalibrated by Picard (version 2.21.8) and GenomeAnalysisToolkit (GATK4.1.3). SNPs were annotated by SnpEff (LGPLv3), and the SNPs which exclude sites of 0.1 missing rate and minQ value greater than 30 and minor allele frequencies (MAF) lower than 0.05 and minimum mean depth value greater than 8 were regarded as high-quality SNPs. Non-synonymous diversity per non-synonymous site (πN) and synonymous diversity per synonymous site (πS) were calculated for potential selection of *B. stacei* and *B. hybridum* using SNPGenie (v 1.0) (Nelson et al., 2015).

## Data availability

Data of this work are available within the paper and its Supplementary Information. The raw reads for the RNA sequences and whole genome resequencing are available from NCBI BioProjects (PRJNA628850) and (PRJNA688592).

## Acknowledgements

This work is funded by the Australian Research Council (FT210100366), the National Natural Science Foundation of China (31620103912, 31571599, and 31571578). PC was funded by the Spanish Ministry of Science and Innovation TED2021-131073B-I00 and PID2022-140074NB-100 grants. We thank Prof. Kexin Li and Xiaoying Song for providing the seeds.

## Author contribution

Z-HC and EN conceived and designed the research. YW, GC and Z-JY performed the experiment. YW conducted data analyses with support from GC, ZH, FX. YW, PC and Z-HC wrote the manuscript with contribution from all authors.

## Supplementary Materials

**Figure S1.** Top 10 GO terms for ortho-homeolog genes with the ratio of nonsynonymous (Ka) to synonymous (Ks) substitution rates greater than 1 between *B. hybridum* and *B. stacei*. BP, Biological Processes; MF, Molecular Functions. GeneRatio, number of observed divided by the number of expected genes from each GO or KEGG category in the gene list.

**Figure S2.** Physiological and stomatal traits of each *Brachypodium* accession collected from African slope (AS; *B. hybridum*) and European slope (ES; *B. stacei*) under well water condition (Control) and drought. Net CO_2_ assimilation (A), stomatal conductance (B), Photosynthetic water use efficiency (*WUEi*) (C), Aperture area (D).

**Figure S3.** Global transcriptome analysis of drought response in *Brachypodium hybridum* (AS) and *Brachypodium stacei* (ES). (A-D) RNA sequencing principal component (PC) analyses revealed that approximately 58.6%, 48%, 38%, 47.5% percent of variation among accessions obtained for this study was explained by the first two principal components. AS and ES, BhS and BhD, Bhs and Bs, BhD and Bs, are distinguished in the first PC, well water condition or drought treatment can be distinguished in the second PC.

**Figure S4.** Top 10 GO terms for Bhs/Bhd dominant homeologs. BP, Biological Processes; CC, Cellular Components, MF, Molecular Functions. Gene Ratio, number of observed divided by the number of expected genes from each GO or KEGG category in the gene list.

**Figure S5.** GO and KEGG enrichment analysis of highly connected genes (hub genes) in *B. hybridum* (A) *and B. stacei* (B). Gene Ratio, number of observed divided by the number of expected genes from each GO or KEGG category in the gene list.

**Figure S6.** Overview of protein interaction network for WGCNA hub genes in MAPK pathway of *Brachypodium hybridum* (A) and *B. stacei* (B). The protein interaction network involving 143 proteins linked via 558 interactions. Brown, blue, red color represent hub genes of Bh, Bs, and both Bh and Bs, respectively. Shade of label color is decided by DEGs absolute value, genes with grey color are not differently expressed. Solid lines connecting the nodes represent direct interaction and the light dots represent other interactions.

**Figure S7.** Protein interaction network for WGCNA hub genes in MAPK and ABA pathway of *Brachypodium hybridum* (A) and *B. stacei* (B). The protein interaction network involving 143 proteins linked via 558 interactions. Brown, blue, red color represent hub genes of Bh, Bs, and both Bh and Bs, respectively. Shade of label color is decided by DEGs absolute value, genes with grey color are not differently expressed. Solid lines connecting the nodes represent direct interaction and the light dots represent other interactions.

**Figure S8.** Circos diagram for genome resequencing and long terminal repeat (LTR) distribution (A), GO terms for genes with high impact SNP (B) in *B. hybridum and Brachypodium stacei*. The Circos plot illustrates key genomic features across four concentric circles. The outermost circle displays the distribution of Long Terminal Repeat (LTR) retrotransposons on each chromosome, with blue bars representing Copia-domain distribution and orange bars indicating Gypsy-domain distribution. The second circle shows the number of high impact in Bh (blue) and Bs (orange). The third circle colored in red presents the density of single nucleotide polymorphisms (SNPs) across the chromosomes. The innermost circle (green bars) represents the density of Indels across the chromosomes.

**Figure S9.** Top10 GO terms for homeolog genes (Ka/Ks >1) between *Brachypodium hybridum* and *Brachypodium stacei.* Gene Ratio, number of observed divided by the number of expected genes from each GO or KEGG category in the gene list.

**Figure S10.** Homeolog expression bias (HEB) for each Brachypodium accession. Under control conditions, around 500 genes are commonly dominated by both the S and D subgenomes across all accessions, but the number of commonly dominated genes decreases to 178 and 162 for the S and D subgenomes, respectively, in response to drought.

**Table S1.** Synteny blocks between Bs and Bh sub-genomes.

**Table S2.** LTR distribution of Bh and Bs genomes.

**Table S3.** RNAseq and re-sequencing raw reads availability.

**Table S4.** Homeologs between BhS and BhD sub-genomes which shifted their dominant subgenome after drought treatment.

**Table S5.** Annotation for genes with high impact SNP in Bh and Bs.

**Table S6.** Homeologs under positive selection (Ka/Ks > 1) in Bh from both subgenomes, stacei-type subgenome (Bs) and distachyon-type subgenome (Bd).

